# Phylogenomic and comparative genome analysis of the Juglandaceae family reveals black walnut (*Juglans nigra*) as an ancient lineage of the *Rhysocaryon* section

**DOI:** 10.1101/2025.11.25.690472

**Authors:** Aziz Ebrahimi, Samarth Mathur, Mohammad Mehdi Arab, Nicholas LaBonte, Evan Long, J. Grey Monroe, Douglass F. Jacobs

## Abstract

Accelerated climate change causes heatwaves, droughts, and severe freezes, posing threats to tree regeneration and to global biodiversity. Deeper genomic insights are crucial for breeding, conserving genetic resources, guiding management, and building resilience to these threats. *Juglans*, a globally distributed genus of timber and nut-producing trees, still has unresolved species relationships; comprehensive genomic data would clarify adaptation to rapid climate change and inform assisted migration strategies. We analyzed 20 *Juglans* species from North and South America and Asia using whole-genome data, chloroplast genome data, cold-hardiness gene families, and environmental variables related to freezing tolerance. Phylogenomic analyses indicate that black walnuts (*J. nigra*, section *Rhysocaryon*) may have diverged ~28.7 MYA (late Oligocene). *J. nigra* reveals higher nucleotide diversity with more genes than other *Juglans*, whereas *J. hindsii* clusters with Central/South American species, consistent with a more recent divergence within *Rhysocaryon*. As expected, *Juglans* species originating from mild-winter regions are distinct from those in cold-climate regions in genome-wide and cold-hardiness gene analyses, with introgression occurring mainly from temperate into tropical lineages. *Juglans* species from milder or warmer winters show CBF gene deletions, whereas cold-adapted species lack such gene deletions. These results support *J. nigra* as an ancient lineage within *Rhysocaryon,* consistent with subsequent colonization of Central/South America and longer evolutionary histories. Genome analyses further suggest *Rhysocaryon* is ancestral to *Cardicaryon, Trachycaryon,* and *Dioscaryon.* These findings illuminate the demographic history of the *Juglans*, particularly *Rhysocaryon,* and have direct implications for gene conservation, breeding, and assisted migration.

## Introduction

Latitudinal gradients create heterogeneous environments that drive local adaptation in trees. However, the environmental factors influencing adaptive evolution remain poorly understood. Both biotic and abiotic conditions shape species distributions across latitudes, with diversity peaking in the tropics and declining toward temperate regions. Classic and modern theories highlight the role of temperature uniformity in the tropics versus variability in temperate zones as major drivers of diversity and adaptation (Janzen, 1967; Jansson et al., 2013; Kerkhoff et al., 2014; Liang et al., 2022; Leites et al., 2023). Plants at higher latitudes experience broad seasonal variability and freezing stress, requiring physiological adaptations to persist and expand their ranges (Bradshaw, 1965; Schubert et al., 2019).

Freezing tolerance varies within and among species, especially those spanning latitude or altitude gradients (Zhen & Ungerer, 2008; Ebrahimi et al., 2023). Cold stress is a primary environmental factor that limits plant growth and shapes geographic distribution (Sanghera et al., 2011). Temperate and arctic plants undergo cold acclimation, triggered by short days and low autumn temperatures, to survive freezing stress (Thomashow, 1999; Monroe et al., 2016). This process enhances membrane stability and osmotic balance through complex molecular pathways (Thomashow, 1999; Hannah et al., 2006). C-repeat binding factor (CBF) proteins are central to this response, which act as key regulators (Thomashow, 1999). Their variation closely correlates with winter temperatures (Thomashow, 1999; Monroe et al., 2016; Lee et al., 2024), and evidence shows that CBFs improve cold tolerance in crops and woody species. In *Arabidopsis thaliana*, reduced nucleotide diversity at CBF2 indicates a selective sweep, suggesting adaptive regulatory evolution (Lee et al., 2024). Similarly, reduced nucleotide diversity and heterozygosity have been observed in CBF and cold-related genes in tree species such as *Betula fujianensis* and *Acacia koa, which were* collected across different latitudes and altitudes (Ebrahimi et al., 2023; Zhuang et al., 2025). Conversely, relaxed selection in milder or warmer climates can reduce CBF function, leading to decreased freezing tolerance in model plant and tree species (Zhen & Ungerer, 2008; Monroe et al., 2016; Ebrahimi et al., 2023; Zhuang et al., 2025). Broader research links the diversification of transcription factor families to the evolution of stress responses across angiosperms (Lehti-Shiu et al., 2015). Genomic studies reveal that structural variants in CBF-dependent genes significantly influence tree adaptation to cold stress (Zhuang et al., 2025). However, connecting genes to adaptive traits during tropical–temperate transitions remains a significant challenge (Wiens & Donoghue, 2004).

The walnut family (Juglandaceae) is a model for studying evolution and climate adaptation. It has ecological and economic importance for species across the Americas and Asia’s tropical, subtropical, and temperate regions (Manchester et al., 2024), and some of the temperate *Juglans* species can tolerate temperatures of −40 to −50 °C (George et al., 1977; Jacobs et al., 2008; Brennan et al., 2021). The family includes ten genera, such as *Juglans*, which is divided into four sections: *Rhysocaryon* (black walnuts, New World), *Cardiocaryon* (Asian butternuts), *Trachycaryon* (*J. cinerea*, North American butternut), and *Dioscaryon* (*J. regia*, Persian walnut) (Manchester, 1989). Fossil evidence shows a long evolutionary history for the *Juglans* species. Middle Eocene fossils from Axel Heiberg Island (above 78°N) indicate that *Rhysocaryon* once occupied much more northerly ranges than it does today (Manchester et al., 2024). Based on fossil data, the lineage likely originated in the mid-latitudes of North America, expanded into higher latitudes across continents by the middle Miocene, and subsequently contracted to mid-latitude regions (Manchester, 1989; Zhang et al., 2022). The earliest well-preserved *Juglans* fruits from the Eocene of North America mark the divergence between the butternuts (*Trachycaryon, Cardiocaryon*) and black walnuts (*Rhysocaryon*) (Scott, 1953; Manchester, 1989). By the Miocene, black walnut fossils were common across North America and later appeared in South America, with occasional finds in Eurasia, suggesting episodic dispersal or preservation and indicating the presence of temperate species (Brown, 1946; Palamarev, 1994; Sakala & Gryc, 2011). Phylogenetic and paleoclimatic studies further suggest that wind-pollinated Juglandaceae diversified in high-latitude regions, becoming highly adapted to cold climates during the Middle Miocene Climatic Optimum (Zhang et al., 2022). Today, *Juglans* ranges roughly from 28°S to 52°N across two continents, making it an important model for genetic and evolutionary studies (Aradhya et al., 2007; Stone et al., 2009; Arab et al., 2020), yet the evolutionary factors underlying cold adaptation and the divergence of these species remain unknown.

Temperature probably limited *Juglans*’ dispersal and speciation by acting as a physiological barrier. After the Last Glacial Maximum, North American *Juglans* expanded from refugia into their current ranges (Manchester, 1989; Brown, 1946; Zhang et al., 2022). Section *Rhysocaryon* is the most species-rich group, with about 16 species. It has a primary diversity hotspot in Mexico, followed by the USA, Central America, and South America (Stanford et al., 2000; Stone et al., 2009). Although temperate *Juglans*, especially *Rhysocaryon*, are common and widespread, many subtropical and tropical relatives have limited ranges and are currently threatened with extinction (IUCN Red List). Relationships within *Rhysocaryon* are still debated; early morphological and molecular studies suggested a Central American origin (Manning, 1957; Fjellström & Parfitt, 1995), whereas newer fossil evidence points to a mid- to high-latitude North American origin (Zhang et al., 2022; Manchester et al., 2024). Genome projects have mainly focused on assemblies for a few *Juglans* species (Stevens et al., 2018), leaving the phylogeny of the *Rhysocaryon* section unresolved and its latitudinal adaptations an open question.

Within *Rhysocaryon*, *J. nigra* (black walnut) holds significant ecological and economic value. It is a widely distributed and more abundant species of this section across eastern North America, between 30-45°N, and is prized for its timber and nuts, making it one of the most common hardwoods within its range (Michler et al., 2007; Ebrahimi et al., 2022). Its broad ecological tolerance likely results from extensive pollen-mediated gene flow and natural selection across various climate zones (Austerlitz et al., 2000). Genomic analyses reveal higher nucleotide diversity in *J. nigra* than in *J. microcarpa* and *J. major*, along with an enrichment of GC/CG motifs. (Stevens et al., 2018; Ebrahimi et al., 2019; Ebrahimi et al., 2020), Furthermore, phylogeographic studies suggest post-glacial recolonization from southern refugia (Williams et al., 2004; Clark et al., 1998), while fossil data indicate the opposite, with more fossil evidence of *Rhysocaryon* found in the mid-latitudes of the northeastern part of the forest compared to the western coast and the southern range in America (Scott, 1953; Manchester, 1989; Zhang et al., 2022). However, the specific genomic mechanisms underlying *J. nigra’s* ability to adapt to different climates and how this species relates to *Rhysocaryon* members remain poorly understood.

Limited genomic resources have long restricted evolutionary studies in the *Rhysocaryon* section. The chromosome-scale assemblies of *J. regia* now provide tools for genome-wide comparisons (Martínez-García, 2016; Marrano et al., 2020). Expanding these resources to *Rhysocaryon* presents an opportunity to investigate how latitude and climate impact genomic diversity and adaptation. In this study, we addressed two questions: (1) What are the evolutionary relationships and patterns of genetic diversity across Juglandaceae, especially within *Rhysocaryon*? (2) How do temperature gradients affect adaptive genetic variation? To answer these, we analyzed the genomes of 20 *Juglans* species across latitudinal gradients. We reconstructed phylogenetic relationships and performed comparative genome analyses using whole-genome, chloroplast, and cold-hardy SNP data. We examined how temperature variation affects diversity and evolution, and assessed climate adaptation through selective sweeps, studies of *Juglans* CBF genes, and heterozygosity analyses.

## Materials and Methods

### *Juglans* species distribution and bioclimate data

*Juglans* species are winter-deciduous, growing in an extended latitudinal distribution across heterogeneous environments and different continental zones. Minimum temperatures in the habitats of *Juglans* species vary widely (**Supplementary Figures 1 and 2**). Based on the GPS coordinates for each sampled individual, the daily minimum temperature for 33 years (1983-2016) was extracted from the CHIRTS-daily data set by the University of California, Santa Barbara Climate Hazards Center (http://data.chc.ucsb.edu/products/CHIRTSdaily/). Nineteen bioclimatic data points from 1998 to 2010 were downloaded from the CHELSA database (https://chelsa-climate.org/bioclim/). The minimum temperature (T_min_) differences across latitude and bioclimate data were analyzed using the tidyverse and ggplot R packages.

### Genome alignment and gene annotation

Paired-end genome data of 210 individuals of *the Juglans* and *Carya* species were downloaded for the NCBI database using the Illumina*®* NovaSeq™ sequencing system (Hu et al., 2017; Stevens et al., 2018; Ebrahimi et al., 2019; Zhang et al., 2019; Zhou et al., 2021). We checked the raw data and removed adapter sequences from each read using Trimmomatic v. 0.36 (Bolger et al., 2014). Low-quality sequences (*Phred <20*) and reads smaller than 30 bp were removed before any downstream analysis. To assemble the whole genome and chloroplast, we applied Burrow-Wheeler Aligner (BWA) to align the paired-end data to the *J. regia* reference nuclear and chloroplast genome, Picard tools to remove duplicated regions (Langmead et al., 2009; McKenna et al., 2010; Wysoker et al., 2013; Marrano et al., 2020), and Genome Analysis Toolkit (GATK) to process mapped reads (McKenna et al., 2010). GVCF tools were combined to process each .vcf file, and then vcftools was used to extract the consensus genome as a FASTA file. AUGUSTUS, the consensus genome represented as a FASTA file, was utilized for gene annotations (Stanke et al., 2006).

To assemble the cold-hardy gene family, the predicted genes in FASTA format were aligned to the UniProt database using DIAMOND in Blastp default mode with ‘DREBs-like’ or ‘CBF proteins’ as a keyword (Buchfink et al. 2015). The cold-hardy genes were converted to .bed format using ‘grep’ command. Then, the VCF tools (http://vcftools.sourceforge.net/) were used to realign .bed format to the .vcf file of the whole-genome, followed by ‘--recode’ command to obtain the SNPs (Single Nucleotide Polymorphisms) related to CBF genes.

### Phylogeny and comparative genome analysis

Phylogenetic trees were constructed using the Assembly and Alignment-Free (AAF) algorithm (Fan et al., 2015) for 210 samples, including 20 *Juglans* and 18 *Carya* species from around the world. In addition, we used SNPs identified across the three datasets (reference-based whole-genome assemblies, chloroplast genomes, and CBF genes) for phylogenetic analysis. (**Figure 1 A, B**; **Supplementary Figure 3 A, B, C**). The combined .vcf file was first filtered with VCF-tools using: ‘--maf 0.05, --minQ 40, and --max-missing 0.8 to remove low-quality SNPs. The high-quality SNPs were converted to HapMap format using TASSEL software (Bradbury et al., 2007). A similarity matrix was calculated using the neighbor-joining method (Nei, 1972), and the newic format was visualized for the phylogenetic trees using <u>itol.embl.de.</u> Divergence-time estimation dated for *Juglans* species using RelTime in MEGA software (Tamura et al., 2018) on the ML Newick tree with fossil MRCA calibrations, yielding an ultrametric chronogram with 95% Cis.

**Figure 1.**
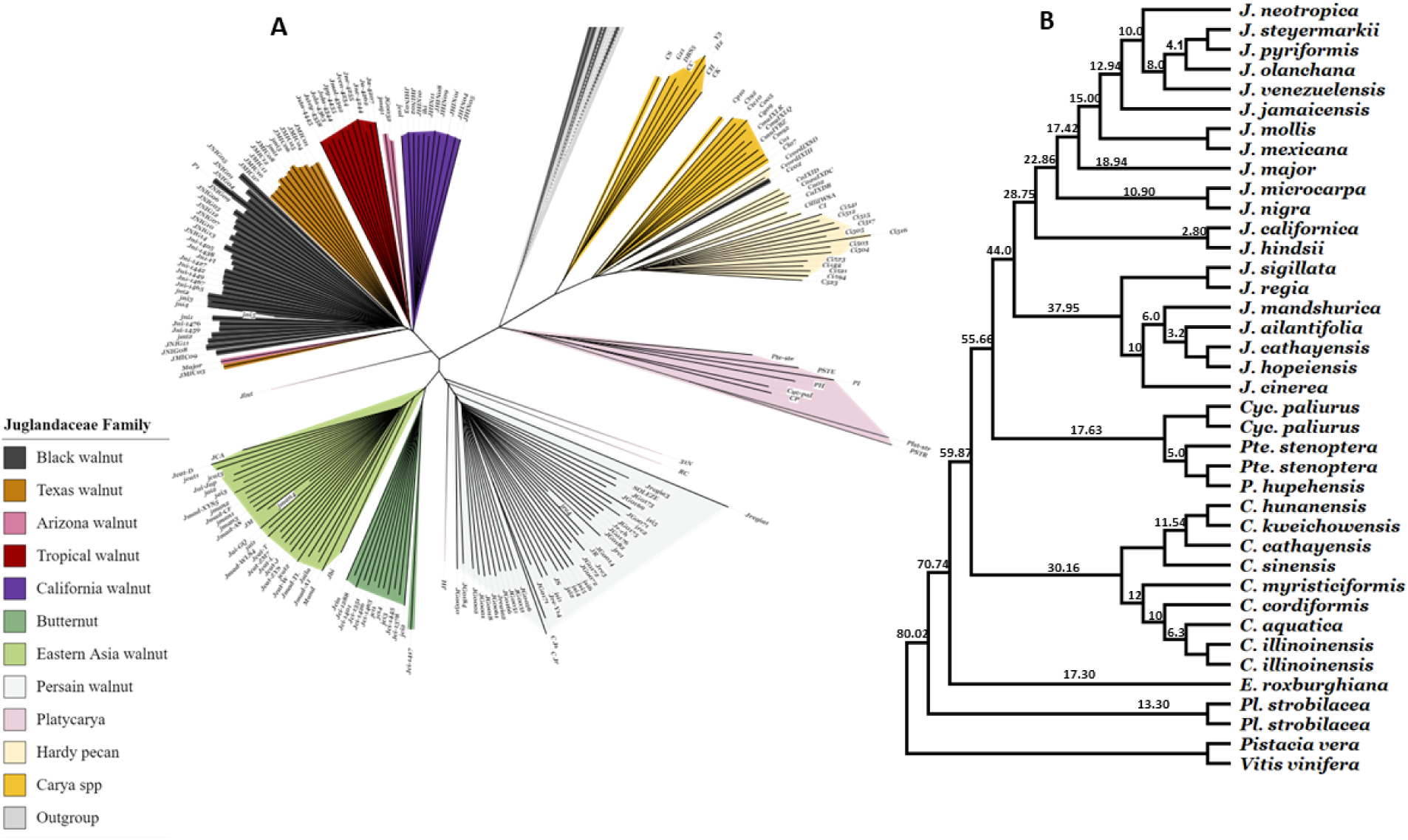
Phylogeny and comparative genome analysis of the Juglandaceae family, including 21 *Juglans* and 18 *Carya* species worldwide. (**A**) Based on the whole genome analysis (n=210); (**B**) Based on the whole genome analysis (n=38). The numbers on the branches indicate divergence times (million years ago) among *Juglans* species.

For comparative genome analysis, orthologous gene families were identified using *OrthoFinder* v2.5.4 (Emms & Kelly, 2019) across the *Juglans* genome. A species tree was constructed using the STAG method (Emms & Kelly, 2018), with node support values representing the proportions of orthogroup trees. Orthogroups limited to a single species or containing more than 100 genes were excluded from analysis in CAFE v5 (Mendes et al., 2021) to detect significant gene family expansion or contraction (*P < 0.01*). The longest gene in each orthogroup was annotated using EnTAP, and retroelement families were removed according to the method described by Guzman-Torres et al. (2024). Orthogroups found in all species were defined as core, those in multiple but not all species as variable, and those missing from a single species as lost; groups unique to one species were considered lineage specific. Classification was validated against the *OrthoFinder* species tree. The upset plot was visualized using the *R ComplexUpset* library (**Figure 2**).

**Figure 2.**
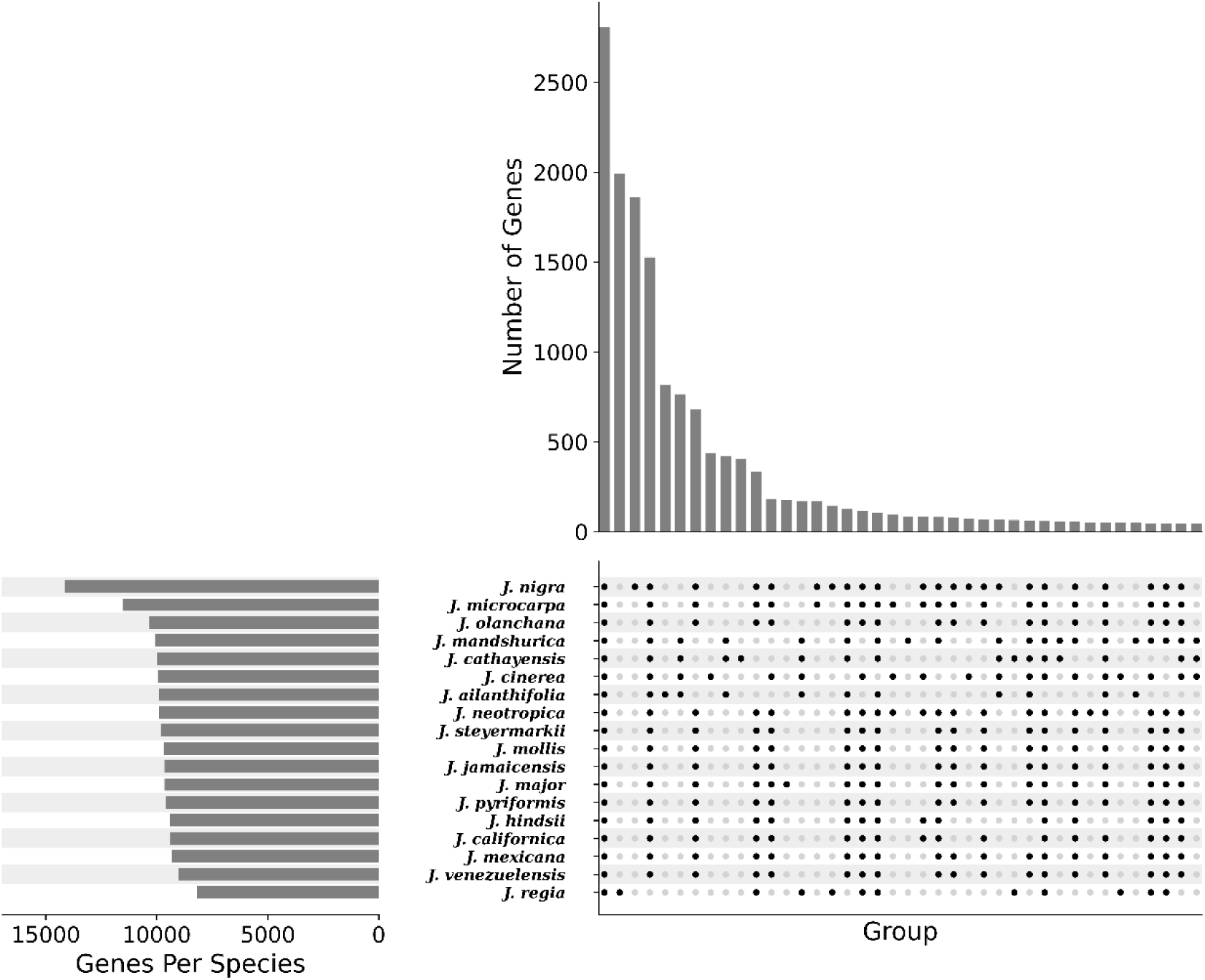
Upset plot of the *Juglans* comparative genome analysis. Intersection bars summarize presence–absence variation (PAV) of gene families across genomes (connected dots). The left bars show the set of genes per species.

### Structure analysis

Plink software was used to convert the .vcf file to the PLINK format for the fast-STRUCTURE software (Raj et al., 2014). Fast-STRUCTURE software was set up for 10 independent runs (K = 2 to 10) with a minimum allele frequency (MAF) of 0.05 and a tolerance of 10^ (−6) for convergence of the log-likelihood difference in 50 iterations, with a maximum of 50,000 iterations. The model complexity (determining which K is better) was calculated using a Python program that selects the optimal K based on the criteria described by Evanno et al. (2005). The mean Q obtained from Fast-STRUCTURE results was used as input to visualize the plots in the R program (Team, 2013) (**Figure 3**).

**Figure 3.**
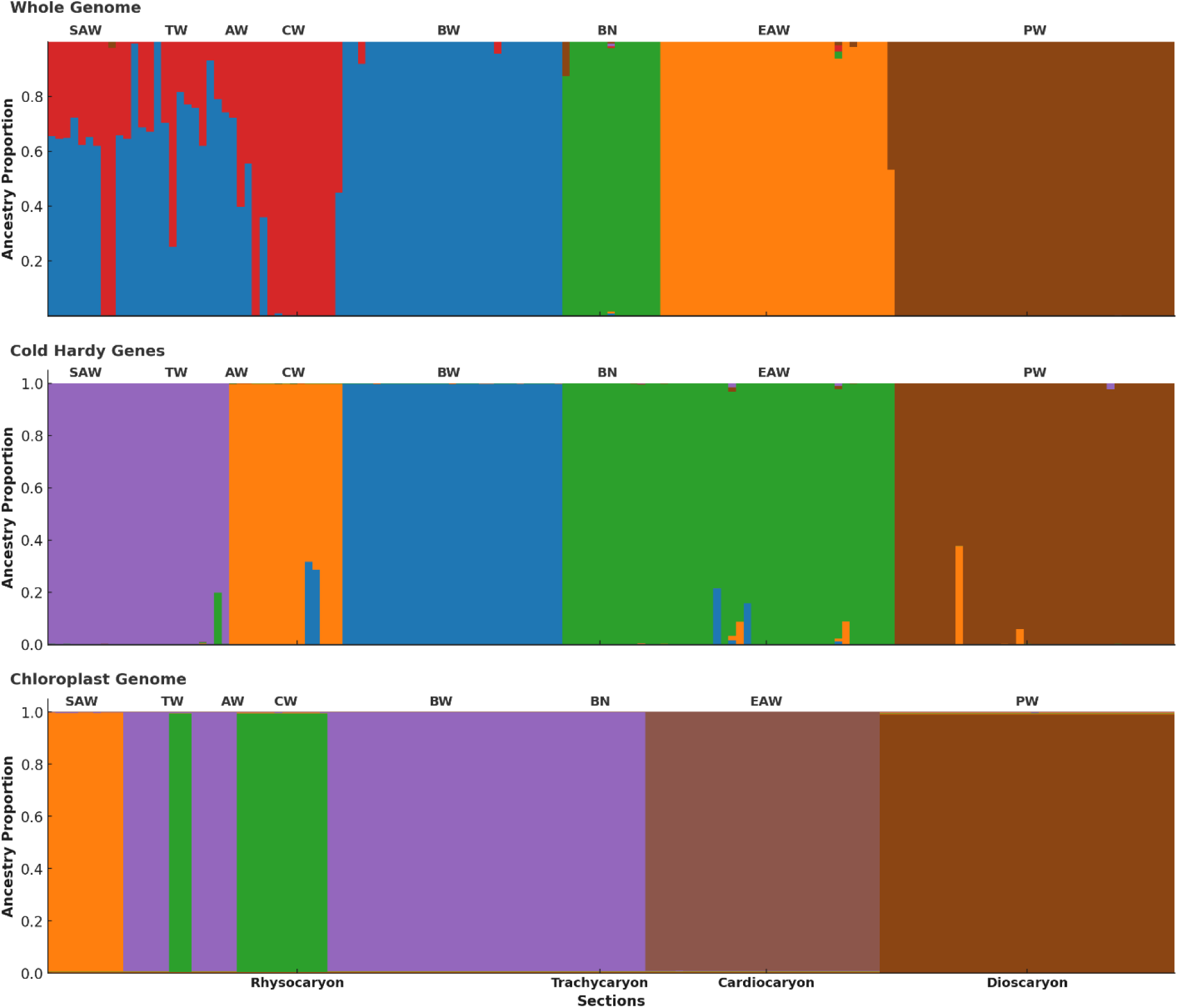
Population structure of 21 *Juglans* spp. across the world, based on the best K numbers. **(Whole Genome)** Analyzed based on SNPs of the nuclear genome (Cold Hardy Genes), SNPs of cold-hardy genes, and (**Chloroplast Genome**) SNPs of the chloroplast genome. SAW stands for Southern American Walnut (including Central America); TW: Texas Walnut; AW: Arizona Walnut; CW: California Walnut; BW: Black Walnut; BN: Butternut; EAW: Eastern Asian Walnut; PW: Persian Walnut.

### Selective sweep of whole genome and cold hardy genes in *Juglans* species

In this study, we used selective sweep analysis of whole genomes and cold hardy genes as separate groups: Japanese walnut (*J. ailantifolia*); Chinese walnut (*J. cathayensis*); Manchurian walnut (*J. mandshurica*) Iron walnut (*J. sigillata*); Black walnut (*J. nigra*); butternut (*J. cinerea*); Arizona walnut (*J. major*); and Persian walnut (*J. regia*); California walnut (*J. hindsii*); Texas walnut (*J. microcarpa*); and tropical walnuts (including all the species that grow in Central and South America and the Caribbean). Tajima’s *D* coefficient was used to detect genomic regions under selection, and nucleotide diversity (π) for each genome pool was calculated with VCF tools and Perl scripts described in LaBonte et al., 2018. A negative Tajima’s *D* value and low values for nucleotide diversity (π) for a predicted gene among *Juglans* species were considered evidence for gene selection during adaptation (**Supplementary Figures 4 and 5**). Vcftools was used to calculate observed heterozygosity using all the .vcf files (representing all individuals) as inputs. To calculate observed heterozygosity for each individual, the number of observed homozygous SNPs for each individual was divided by the total number of SNPs for that individual. (N_SNP-O-Hem)/N_SNP (**Figure 4**). For ABBA-BABBA analysis, we followed the framework of Malinsky et al. (2021) implemented in *Dsuite v0.5 r49*. A species assignment file was generated by querying sample IDs from the filtered VCF and mapping them to species labels of *J. nigra. J. hindsii*, *J. microcarpa*, and southern American *Juglans* species, and the outgroup was *J. cathaysensis*.

**Figure 4.**
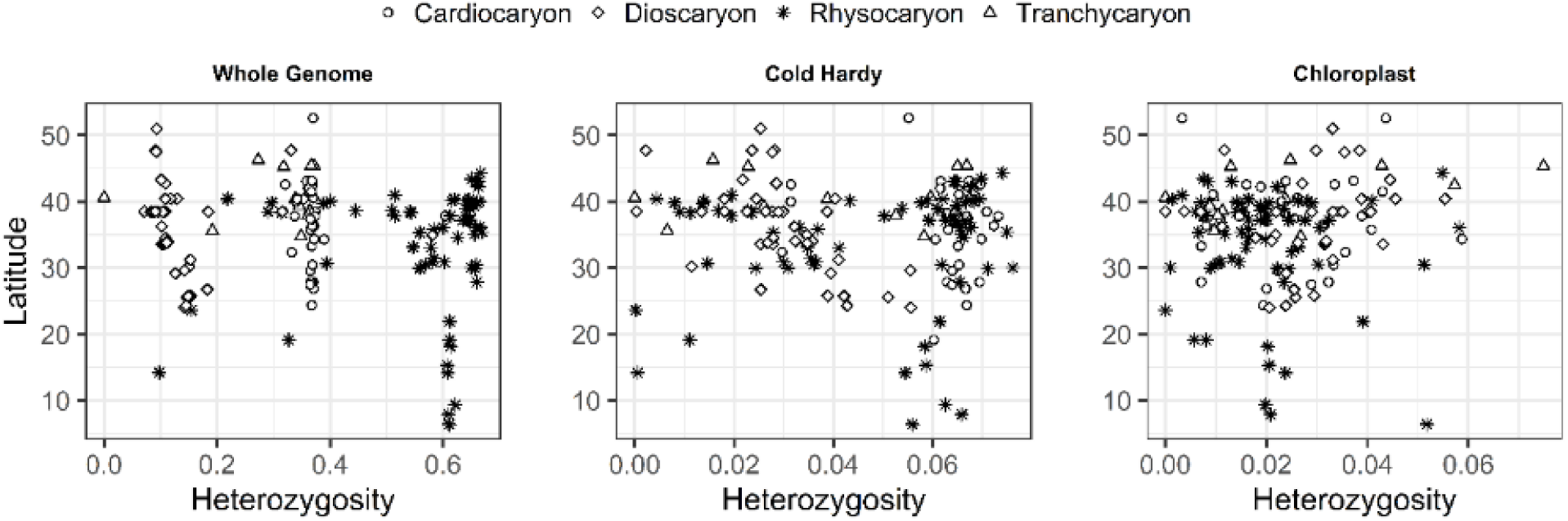
Relationship between latitude and genetic diversity using observed heterozygosity of various *Juglans* species across latitudes based on the whole genome (A), chloroplast (B), and cold-hardy genes (C).

### Climatic and evolutionary insights into *Juglans* CBF genes through protein conservation

The sequences for CBF1, CBF2, and CBF3 were extracted using BLASTn, with the whole genome of each individual as the subject and three references as the query (JX875914.1, JrCBF2, and XM_041146824.1). We analyzed three *Juglans CBF* gene families (*CBF1*, *CBF2*, *and CBF3)* by identifying the most extended open reading frames (ORFs) from curated FASTA files, retaining sequences that began with methionine and ended at the first stop codon. Samples, including geographic coordinates, were merged with 19 bioclimatic variables from WorldClim v2.1, and climatic variation was summarized using principal component analysis (PCA). Protein sequences were aligned using ClustalW, and the consensus was calculated as the proportion of residues matching the most common amino acid at each position (**Figure 5**). Pairwise Hamming distances were computed on the alignments and used for principal component analysis (PCA) to examine sequence divergence. We assessed the associations between climate and protein conservation using logistic regression and Pearson and Spearman correlations to identify climate variables that predict functional sequence loss (**Figure 6**). Visualizations included amino acid alignments, protein length plots, sequence principal component analysis (PCA), and climate-consensus correlation plots, which were all generated in R using the ggplot2 package.

**Figure 5.**
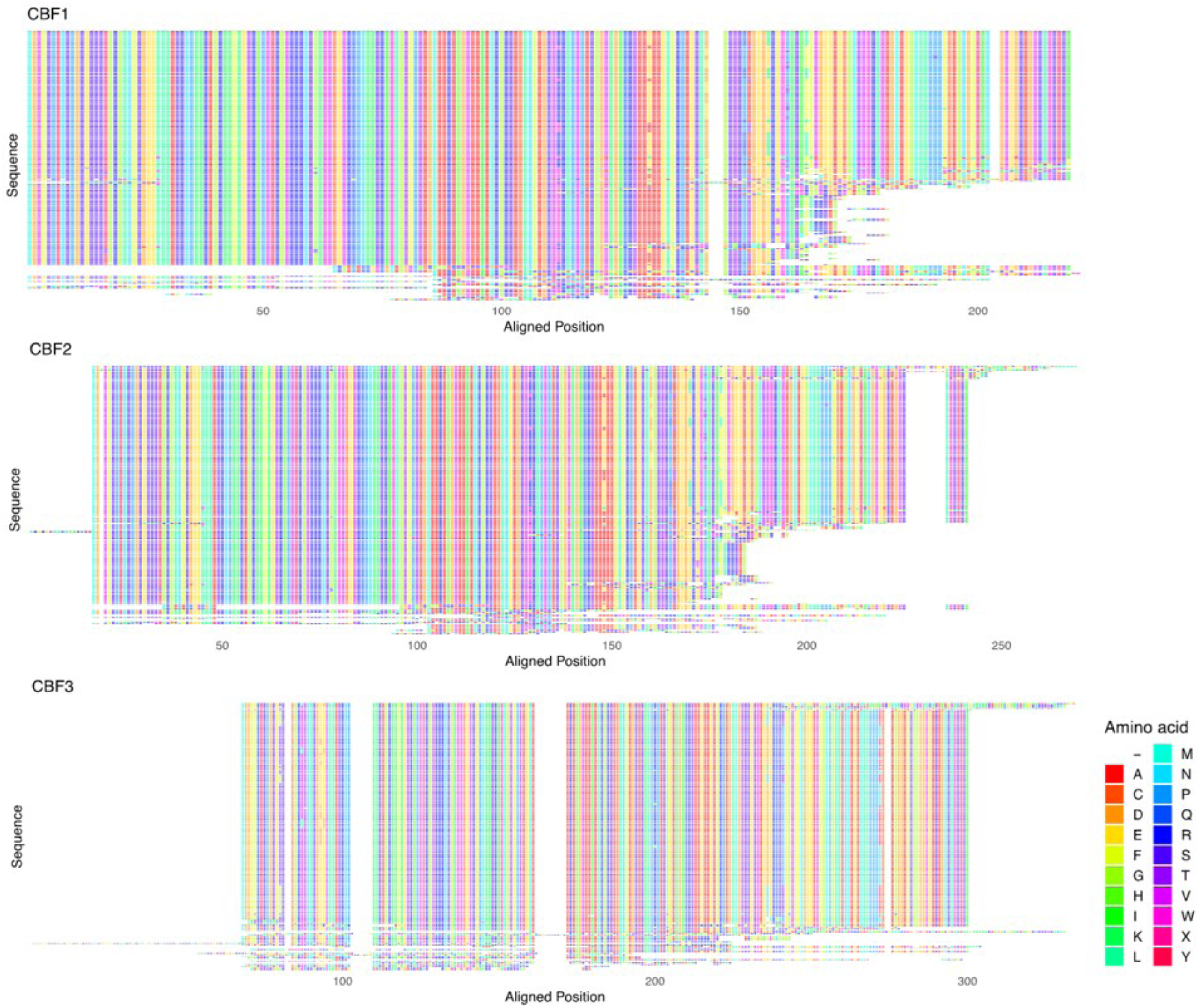
Aligned CBF protein sequences predicted from DNA sequence data. Rows represent individual samples of various *Juglans* species.

**Figure 6.**
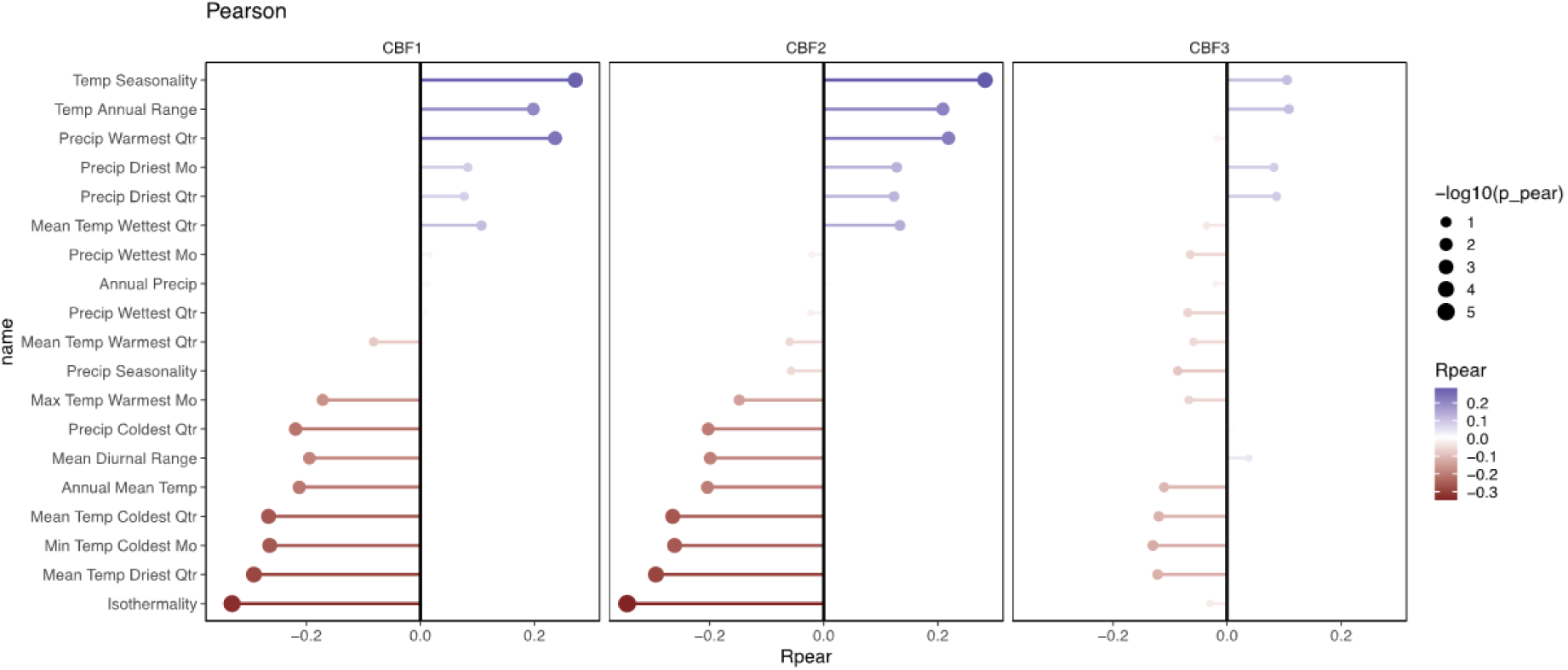
Correlations (Pearson) between CBF sequence (% identity to consensus sequence) and climate of origin. CBF with extreme variation, exemplified by putative loss-of-function variants, is associated with warmer environments, particularly warmer winters.

## Results

### Phylogenomic and comparative genome analysis

Phylogenomic analysis recovered *Juglans* clades that largely mirror geography (**Figure 1A, B**; **Supplementary Figure 3**). A key exception is *J. cinerea*, native to eastern North America, which clusters with East Asian taxa. Section *Rhysocaryon* originated ~28 Ma (Oligocene); *J. nigra* and *J. microcarpa* diverged ~18 Ma. South American species plus *J. major* group with *J. microcarpa* and *J. nigra* with divergences ~22–8 MYA. Unexpectedly, the temperate North American *J. hindsii* and *J. californica* form a clade closer to South American *Juglans*, diverging ~10 MYA (**Figure 1 B**). Chloroplast SNPs place *J. cinerea* near *J. nigra* and *J. microcarpa*, while *J. hindsii* chloroplasts are more similar to South and Central American species than to *J. nigra* and *J. microcarpa*. Cold-hardiness SNPs indicate that tropical species are genetically closer to *J. hindsii*, despite its temperate nature, consistent with a relatively recent origin within *Rhysocaryon* (**Supplementary Figure 3 C**).

Comparative genomics reveals that the number of genes per species ranges from approximately 8,000 to 14,000 (**Figure 2**). Rhysocaryon generally exceeds *Cardiocaryon*, *Trachycaryon*, and *Dioscaryon* in gene counts. Within *Rhysocaryon*, *J. nigra* (followed by *J. microcarpa*) possesses some of the highest gene counts, whereas *J. hindsii* and *J. californica* display fewer genes than other temperate species like *J. nigra*, *J. microcarpa*, and *J. major*, as well as several tropical Central and South American species. *J. cinerea* has the same number of genes as *J. ailantifolia*, while *J. mandshurica* and *J. cathayensis* surpass *J. ailantifolia*. Domesticated *J. regia* has fewer genes compared to other *Juglans* species. These patterns suggest that *Rhysocaryon* is an older, more diverse lineage compared to other sections. *J. hindsii* is a relatively recent lineage, and *J. nigra* possesses a broader gene repertoire within the section, possessing more genes overall among all *Juglans* species worldwide.

### Structure analysis

Structure analysis based on SNPs from whole genomes, chloroplasts, and CBF genes grouped the *Juglans* species into six distinct clusters (**Figure 3**). Within the *Rhysocaryon* section, whole-genome and CBF gene analyses separated it into four subgroups, whereas the chloroplast genome divided it into three groups (**Figure 3**). Whole-genome analyses showed significant gene flow between *J. nigra*, *J. microcarpa*, and southern American walnuts, while *J. hindsii* formed a separate cluster from the rest of the *Rhysocaryon* section (**Figure 3**). Interestingly, southern American species, along with *J. microcarpa* and *J. major*, showed significant gene flow with *J. hindsii*, which is distinct from *J. nigra*. Butternut, *J. cinerea* grouped with *J. nigra* and *J. microcarpa* in the chloroplast-based analysis, yet they did not group with the rest of the *Rhysocaryon* members. Furthermore, ABBA–BABA analyses revealed significant introgression from *J. nigra* into both South American *Juglans* species as well as *J. microcarpa* rather than *J. hindsii*.

### Selective sweep of the whole genome and cold-hardy genes

Selective sweeps were analyzed using SNPs from entire genomes and cold-hardy genes to estimate heterozygosity, *Tajima’s* D, and nucleotide diversity across *Juglans* species (**Figure 4**, **Supplementary Figure 4**, **5**). Species from the American *Rhysocaryon* section had greater heterozygosity than those from Asia (**Figure 4**). Notably, *J. nigra*, found above 35-42°N, showed higher heterozygosity than lower-latitude species such as *J. microcarpa*, southern American walnuts, and *J. hindsii*, a temperate species (**Figure 4**). Cold-hardy gene heterozygosity clustered temperate North American and East Asian temperate species, suggesting convergent adaptation to similar climates. No clear patterns of chloroplast heterozygosity were observed among continents. Whole-genome analysis showed *J. nigra* with *Tajima’s* D close to zero, while *J. regia* had a negative skew, indicating purifying selection. Most other species displayed neutral or positive *Tajima’s* D, reflecting balancing or positive selection. For cold-hardy genes, *J. regia* and *J. cathayensis* showed strongly negative *Tajima’s* D, which reduced genetic variation, whereas *J. microcarpa* and *J. hindsii* showed slight negative skew with non-normal distributions. In contrast, *J. mandshurica* exhibited positive skew, likely due to the harsh winter conditions in its habitat. Nucleotide diversity for both whole genomes and cold-hardy genes was lowest in *J. regia*, *J. hindsii*, *J. sigillata*, and tropical walnuts, and highest in *J. nigra* and *J. cinerea*.

### Climate-driven CBF gene divergence in *Juglans*

Comparative analysis of *Juglans* CBF1, CBF2, and CBF3 proteins revealed distinct conservation patterns (**Figure 5**). CBF1 and CBF2 maintain strong N-terminal conservation (~1–100 aa and ~1–120 aa, respectively), preserving the AP2 domain. In contrast, their mid-to-C-terminal regions show gaps and truncations in species from warmer or milder winter climates, such as South and Central American walnuts, *J. microcarpa* from Texas, and *J. hindsii* from California, indicating relaxed selection or partial loss of function. CBF3, the longest (~310 aa), displays a modular structure with three conserved blocks separated by large indels, suggesting possible exon variation or sub-functionalization. Pearson correlation analyses between CBF protein conservation and 19 bioclimatic variables (**Figure 6**) revealed strong negative correlations for CBF1 and CBF2 with annual mean temperature, minimum temperature of the coldest month, and isothermality, while showing positive correlations with temperature seasonality and annual temperature range. These patterns suggest that colder, more thermally variable climates favor greater conservation of CBF1 and CBF2, whereas warmer, more stable climates encourage divergence and potential loss of function of alleles. Conversely, CBF3 showed weaker, mostly non-significant climatic associations, likely due to its modular structure or different selective pressures. Overall, these findings suggest that CBF1 and CBF2 are closely associated with cold adaptation and demonstrate relaxed selection in warmer regions, while CBF3 evolves under different constraints. This highlights the crucial role of CBF1 and CBF2 in freezing tolerance, providing insights into the evolutionary mechanisms that shape climate adaptation in *Juglans*.

## Discussion

### Phylogenetic, structural, and comparative genome analyses

Previous studies have investigated *Juglans’* phylogeny using morphology, molecular markers, and fossil evidence; however, the results have often been incongruent. For example, *ITS* and *matK* markers produced poor resolution within the *Rhysocaryon* section, with morphological traits dividing species into three groups, whereas molecular data placed them into two (Dode, 1906; Stanford et al., 2000; Aradhya et al., 2007). Such inconsistencies reflect the limitations of morphology and a restricted number of markers, many of which evolve too slowly or are influenced by environmental plasticity (Rieseberg & Soltis, 1991; Stanford et al., 2000; Aradhya et al., 2007). Whole-genome approaches, including our study, offer significantly greater power to resolve these relationships, especially within the *Rhysocaryon* section. Our genome-scale phylogeny reveals that most *Juglans* species cluster by geography, except for *J. cinerea*, which groups with eastern Asian species, consistent with earlier studies (Stanford et al., 2000; Zhou et al., 2021). Chloroplast data instead place *J. cinerea* with only *J. nigra* and *J. microcarpa,* not sorting it with the rest of the *Rhysocaryon section,* as they grow in the same habitat range, supporting the hypothesis of chloroplast capture or introgression. Similar organellar capture events are well documented in *Salix* and *Quercus* (Rieseberg & Soltis, 1991), suggesting that hybridization has played a recurrent role in shaping *Juglans* lineages. Fossil evidence corroborates this view, showing that black walnut and butternut lineages diversified during the mid-Eocene, with Arctic fossils confirming that major sections were established by this time (Hills et al., 1974; Aradhya et al., 2007; Manchester et al., 2024).

Our analyses indicate that *Rhysocaryon* is an ancient, species-rich lineage with higher gene counts than *Cardiocaryon*, *Trachycaryon*, or *Dioscaryon*. Within *Rhysocaryon*, *J. nigra* stands out as an ancestral lineage with many genes, whereas *J. hindsii*, which has fewer genes, appears as the earliest divergence and clusters with tropical walnuts rather than temperate *J. nigra* and *J. microcarpa*, consistent with a previous study that mentioned distinctive pollen morphology and limited fossil evidence of *J. hindsii* compared to *J. nigra* (Scott, 1953; Stanford et al., 2000). Contrary to the hypothesis of a Central American origin from *J. olanchana* with northward expansion (Manning, 1957; Fjellström & Parfitt, 1995), our structure and phylogenomic results support diversification in temperate North America in the late Oligocene (~28 Ma), followed by southward expansion into tropical regions and the subsequent evolution of *J. hindsii*, a temperate-to-tropical route that contraries the usual pattern for other woody species (Janzen, 1967; Kerkhoff et al., 2014; Liang et al., 2022). Miocene–Pliocene events further shaped diversification: the Central American land bridge enabled southward migration, while rapid mountain uplift and climatic shifts fragmented habitats and restricted gene flow, especially among Central and South American species such as *J. neotropica* and *J. australis* (Stone et al., 2009; Zhang et al., 2022). Molecular phylogenies and fossils place Caribbean endemics (*J. jamaicensis*, *J. venezuelensis*) in the mid-Pliocene (Graham, 1999; Manos et al., 2007), and fossil-based models trace *Juglans’* origins to mid-latitude North America through high-latitude (Arctic) *Rhysocaryon* and dispersal into Eurasia via Bering and North Atlantic land bridges (Mu et al., 2020; Zhang et al., 2022; Manchester et al., 2024). Overall, in our study, *Rhysocaryon*, particularly mid-latitude *J. nigra*, exhibits higher gene flow, gene counts and heterozygosity than other members of this section and Asian walnuts, consistent with fossil evidence suggesting that mid-latitude North America is the origin of *Juglans* species (Zhang et al., 2022). Recent phylogenetic evidence shows that *Quercus*, the North American oak genus, originated in the temperate zone, migrated south during glacial periods, and underwent major speciation in the mountainous tropics after the climate warmed (Hipp et al., 2020), a pattern similar to what we’ve observed in *Juglans*. *Pinus* offers another potential parallel to *Juglans*, although pines are usually adapted to cold climates. Tropical Mexico has the highest species diversity, and recent phylogenetic studies show that many of these tropical species are evolutionarily young and likely originated from temperate lineages that migrated southward during periods of climate cooling in the early Oligocene, followed by diversification in the Miocene (Jin et al., 2021).

### Parallel cold-adaptation in *Juglans*

Whole-genome analyses showed *Tajima’s* D ≈ 0 in *J. nigra* and (near-zero to slightly negative) in *J. microcarpa*, with mostly slightly positive genome-wide values across *Juglans*, except domesticated *J. regia*, which exhibited a pronounced negative skew (*D = −0.975*) (Braverman et al., 1995). When assessed with negatively skewed distributions at cold-hardiness loci, this pattern indicates domestication bottlenecks and strong human selection for nut traits (Mercuri et al., 2013), concordant with reduced diversity in U.S. Midwest and European populations relative to Asian sources (Ebrahimi et al., 2017; Pollegioni et al., 2017). Species contrasts were marked, mid-latitude *J. nigra* maintained near-zero *Tajima’s* D with high nucleotide diversity, consistent with broad standing variation under cold climates; species from milder regions, *J. microcarpa*, *J. major*, *J. hindsii*, and Central or South American showed slightly negative *Tajima’s* D and partial CBF deletions, whereas *J. nigra* and *J. cinerea* lacked CBF loss. Notably, temperate *J. hindsii* was exceptional, bearing CBF deletions and the lowest nucleotide diversity. In contrast, *J. mandshurica* displayed genome-wide nucleotide diversity *D ≈ 0*, with a slight positive skew at cold-hardy genes, consistent with balancing or diversifying selection during extreme winters. Field evidence of cold hardiness study aligns with these genomic results, at ~40° N in Indiana (2003–2013), eastern Asian *Juglans* and local *J. nigra, J. cinerea* showed no winter injury (Ebrahimi et al., 2020). In contrast, an extreme frost in early February 2014 dropped to −30 °C (~4 h on two consecutive days), causing severe damage in southern-range species, including *J. major*, *J. hindsii*, and *J. regia*, consistent with lower genetic diversity at cold-hardy genes and CBF function loss in our analysis that could lead to maladaptation despite acclimation to site climate of these *Juglans* species for one decade (Ebrahimi et al., 2020). Similar patterns in conifers and *Betula fujianensis* indicate greater cold-hardiness and higher diversity among cold-adapted taxa (Charlesworth et al., 1993; Lu et al., 2019; Zhuang et al., 2025). Overall, the data indicate: (i) strong artificial selection and bottlenecks in *J. regia*; (ii) relaxed selection and partial loss of cold-response genes in warm-climate species; and (iii) maintenance or elevation of diversity in cold-adapted lineages (e.g., *J. nigra, J. mandshurica*). Convergent evidence from Arabidopsis, lettuce (*Lactuca sativa*), *Betula fujianensis*, and *Acacia koa* shows that reduced CBF function can be advantageous in warm winters (Ågren et al., 2013; Oakley et al., 2014; Monroe et al., 2016; Lee et al., 2024; Ebrahimi et al., 2023; Zhuang et al., 2025), situating *Juglans* within a broader, climate-driven parallel evolution of cold-hardiness loci in our study.

### Genetic diversity and evolution of *Juglans* across latitude and continent

Genetic diversity analyses across the *Juglans* range highlighted a strong relationship between genetic diversity and latitude, particularly within the *Rhysocaryon* section, which originated from tropical to temperate regions in the Americas. Nuclear genomes showed higher heterozygosity than chloroplast and cold-hardy genes, particularly across different walnut sections. North American species from mid-latitudes generally displayed greater diversity than those from Asia, reflecting both historical and current processes of isolation by distance and post-glacial colonization. Fossil evidence indicates that Juglandaceae originated in mid-latitude North America, then expanded to the north and east into Eurasia (Manchester, 1989; Zhang et al., 2022; Manchester et al., 2024). The hypothesis of more abundant fossils in mid-latitude North America and the recent discovery of Arctic walnut fossils above 78° N support our observation that mid-latitude and temperate North American species serve as origins of *Juglans* species and possess greater genetic diversity and more genes than other *Juglans.* Species-specific patterns demonstrate how isolation has shaped genetic variation. For example, mid-latitude *J. hindsii* from California displays some of the lowest diversity across nuclear and cold-hardy genes, consistent with structure analyses showing limited gene flow with northeastern American species and supporting its distinct position within *Rhysocaryon*. Our results also indicate that *J. hindsii* is a recent descendant of tropical lineages that subsequently recolonized a temperate region. An alternative scenario is that *J. hindsii* may have experienced a severe, whole-population bottleneck driven by past climate change. To fully evaluate this possibility, additional sampling of California walnut populations, as well as related tropical walnut lineages, will be necessary. *J. regia* also exhibited low genetic diversity and the fewest genes, reflecting a long history of human domestication. Conversely, *J. nigra* maintained higher diversity and more genes, representing an ancient *Rhysocaryon* lineage well-adapted to colder North American climates. It also exceeds that of members of *Cardiocaryon* and *Trachycaryon* in gene numbers and genetic diversity. Recent microfossil evidence of walnuts from Axel Heiberg Island in northern Canada (above 78° N) confirms the hypothesis that, during the Neogene, there was exchange of *Juglans* taxa between North America and Asia and between Europe and Asia, resulting in a broad Miocene distribution of *Juglans* (e.g., section *Cardiocaryon*) (Manchester, 1989; Zhang et al., 2022; Manchester et al., 2024). This microfossil data aligns with our genome data, indicating that *Rhysocaryon* lineages are older than those of other *Juglans* sections.

Cold-response loci provide additional evidence for latitude-linked adaptation. Mid-latitude species from eastern Asia and North America cluster by heterozygosity at cold-hardiness genes, indicating convergent evolution under similar thermal regimes. Field observations of various *Juglans* species also show that these mid-latitude species tolerate severe winters relative to taxa from milder winter climates (Ebrahimi et al., 2020). Fossil records suggest that adaptation to seasonal cold enabled Juglandaceae to expand from mid-latitudes to Canada and Siberia, thereby fostering the diversification and range expansion of *Juglans* (Zhang et al., 2022; Manchester et al., 2024). Progressive cooling and aridification during the late Neogene–Quaternary prompted retreats from higher latitudes, revealing limits to niche evolution (Zhang et al., 2022). Phylogeographic analyses further indicate restricted gene flow between eastern and western North American lineages, but noticeable gene flow between eastern North American and Central and South American species.

## Conclusion

Genome-scale phylogenies and comparative genomics resolve longstanding conflicts in *Juglans*, positioning R*hysocaryon* as the oldest, species-rich lineage. Within this clade, *J. nigra* maintains the highest diversity, gene counts, and adaptive potential, consistent with its role as an old, genetically rich lineage. In contrast, *J. hindsii* has fewer genes and appears as an early-diverging lineage within *Rhysocaryon*, clustering with tropical walnuts rather than with temperate *J. nigra* and *J. microcarpa*. Together with discordance among chloroplast, nuclear, and cold-hardy genes, and with previous evidence of distinctive pollen morphology and limited fossil representation for *J. hindsii* relative *to J. nigra*, these findings support a temperate North American origin in the late Oligocene (~28 Ma), followed by southward expansion into tropical regions and the subsequent evolution of *J. hindsii*. This temperate-to-tropical trajectory contrasts with the more commonly proposed tropical-to-temperate pattern for other woody genera.

Diversity scans and CBF analyses reveal climate-linked selection: domesticated *J. regia* shows signatures of bottlenecks and artificial selection, cold-adapted species retain intact CBF1 and CBF2, whereas warm-climate lineages exhibit truncations or partial loss, consistent with relaxed selection in frost-free environments. This integrated framework, spanning phylogeny, introgression, comparative genomics, and climate-responsive cold-hardiness pathways, clarifies walnut diversification and provides actionable targets for conservation, assisted migration, and breeding under accelerating climate change. Overall, our results suggest that cold tolerance is a basal trait in *Juglans*, retained in the oldest or most conserved lineages (such as *J. nigra*) and secondarily reduced or lost in some more recently evolved species.

## Acknowledgement

I thank James McKenna, Dr. Shaneka Lawson, Dr. Keith Woeste (USDA Forest Service), Dr. Jingjing Liang, and Dr. Yiwei Jiang (Purdue University) for their support. I also gratefully acknowledge funding from the Ross Fellowship, the Hardwood Tree Improvement and Regeneration Center (HTIRC), and the Bisland Foundation. Dr. Sven Nelson (Director of Plant Science, Heliponix, LLC) and Luhanhai Ying (Purdue University) for consulting in data analysis.

---

**Supplementary Figure 1.**
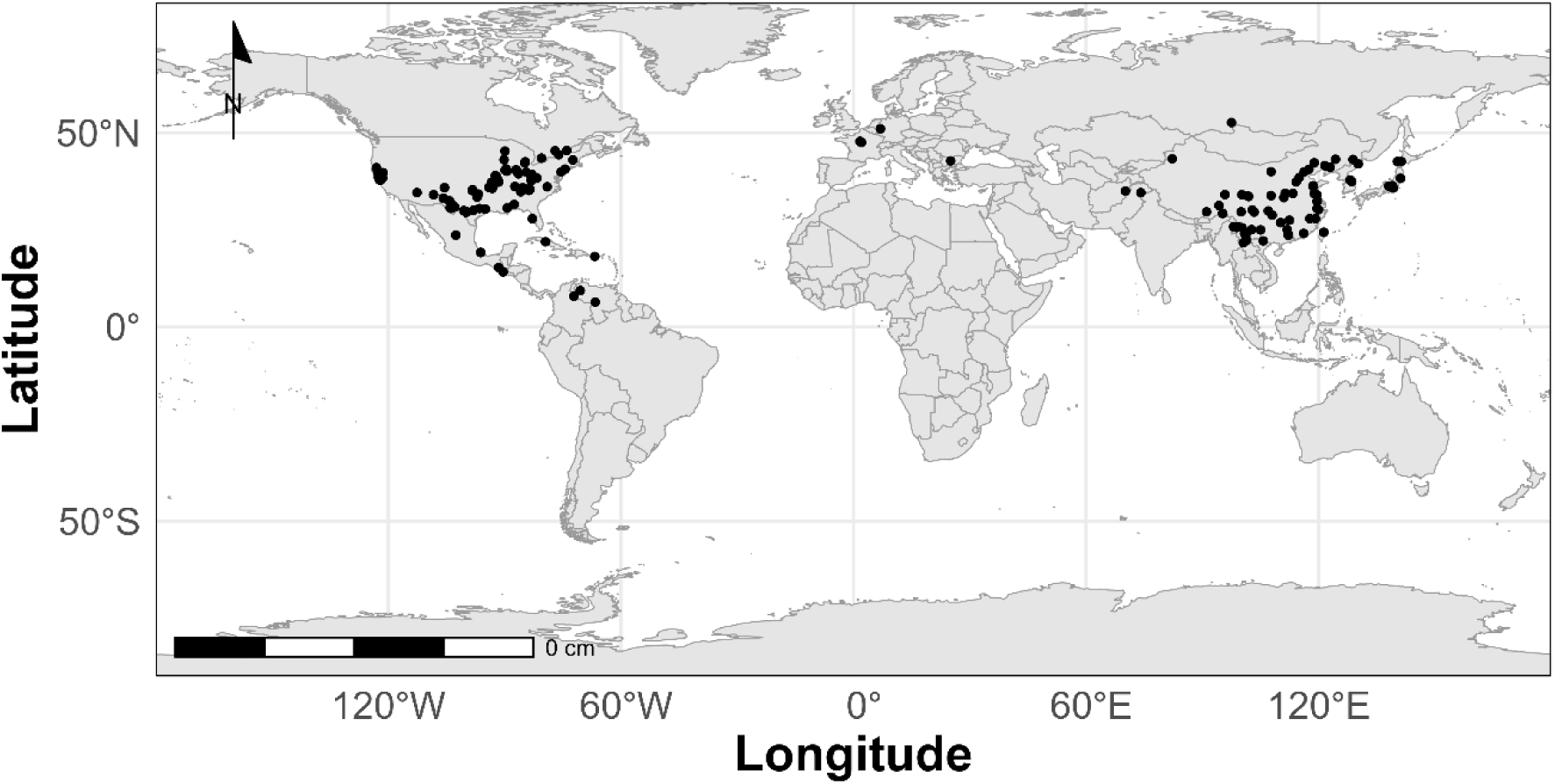
The GPS locations of 21 *Juglans* and 18 *Carya* species (n = 210) used for phylogenomic and comparative genome analysis.

**Supplementary Figure 2:**
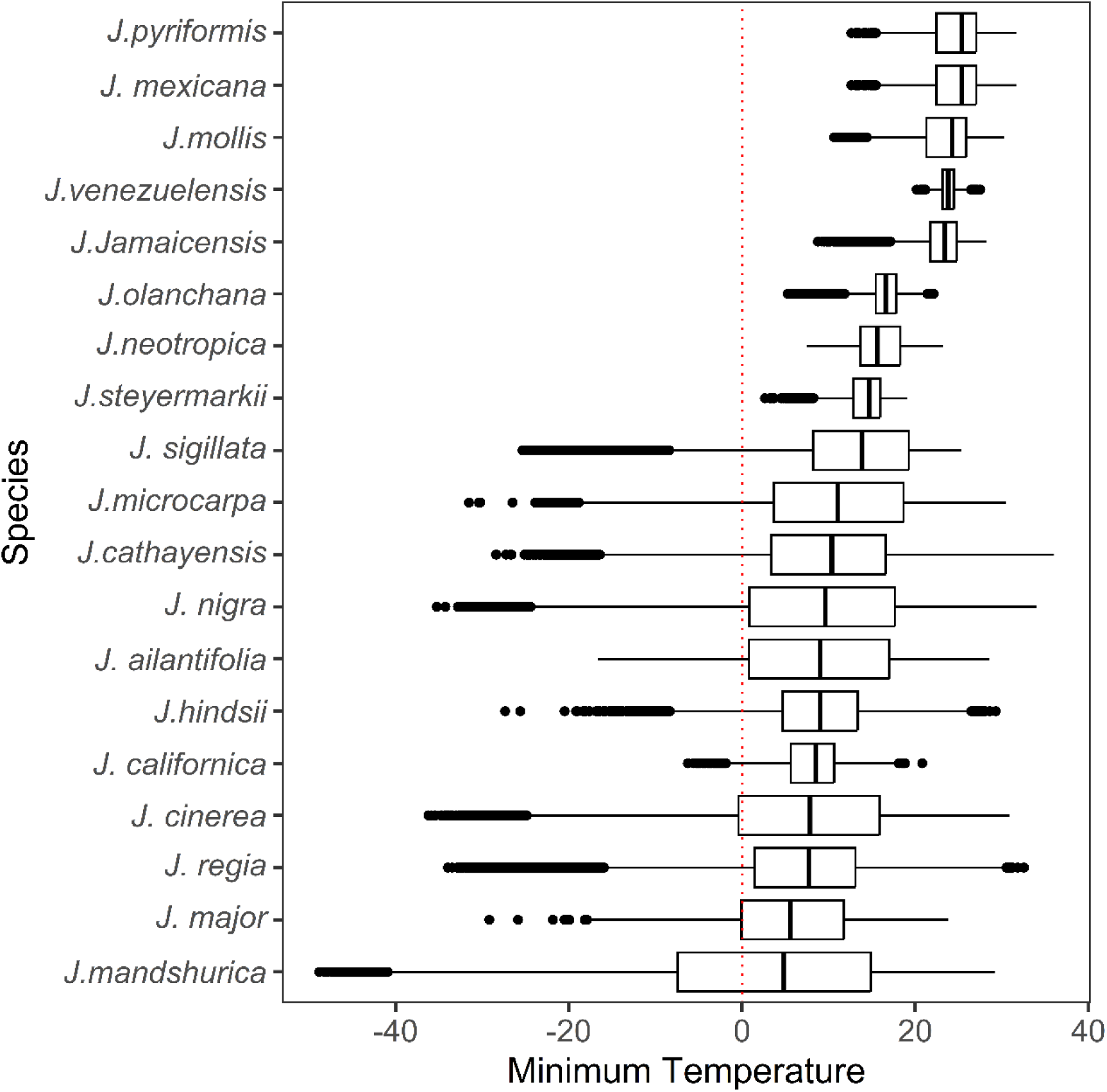
Annual minimum temperature recorded from 1983 to 2016 for different *Juglans* species worldwide.

**Supplementary Figure 3:**
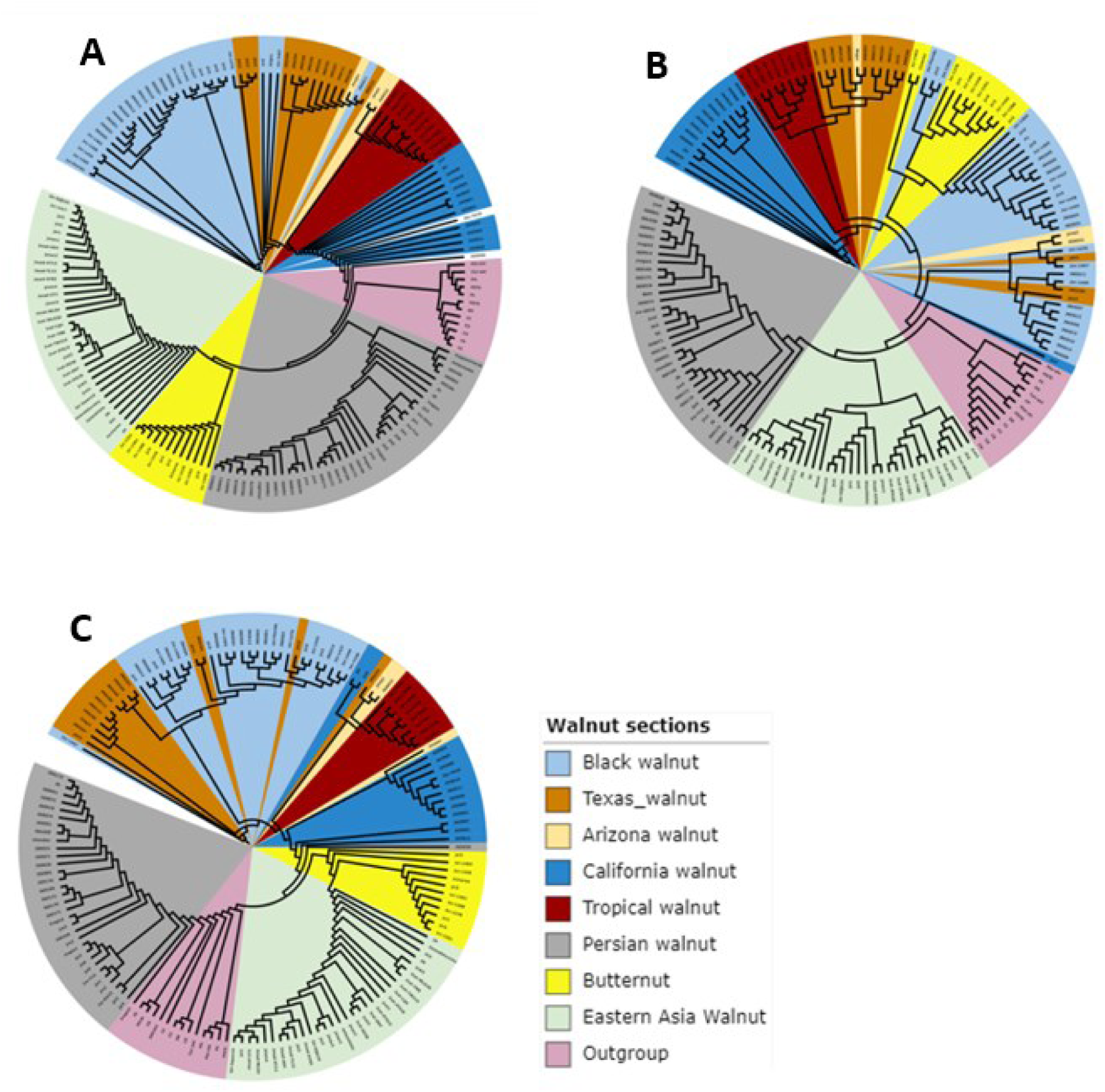
Phylogeny of 21 *Juglans* spp. and *Carya* spp. as an outgroup (n=170) worldwide. **(A)** SNPs of the whole nuclear genome; (B) SNPs of the chloroplast genome; (**C**) SNPs of cold-hardy genes

**Supplementary Figure 4:**
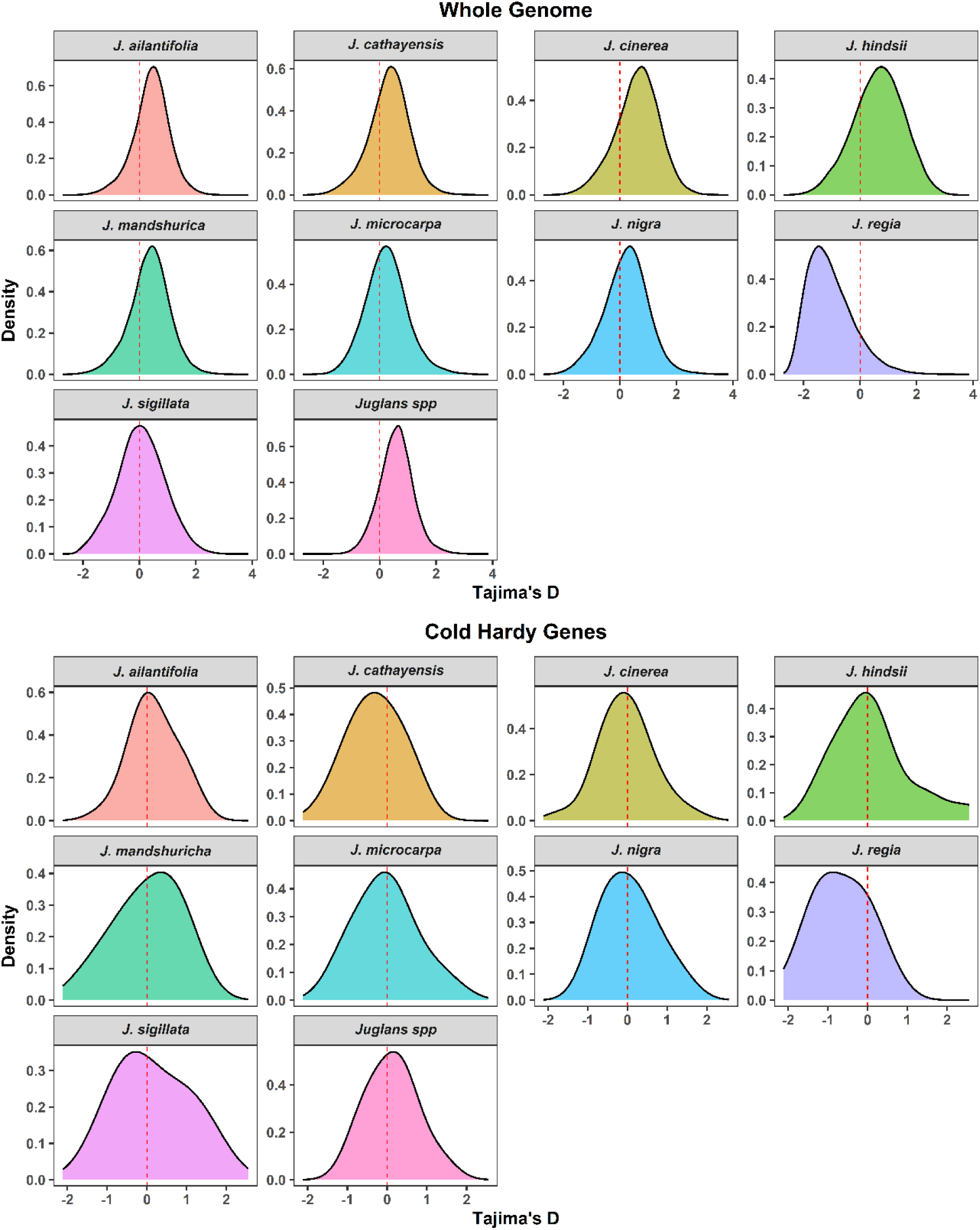
Tajima’s D value of the whole genome and cold-hardy genes across *Juglans* species pool.

**Supplementary Figure 5.**
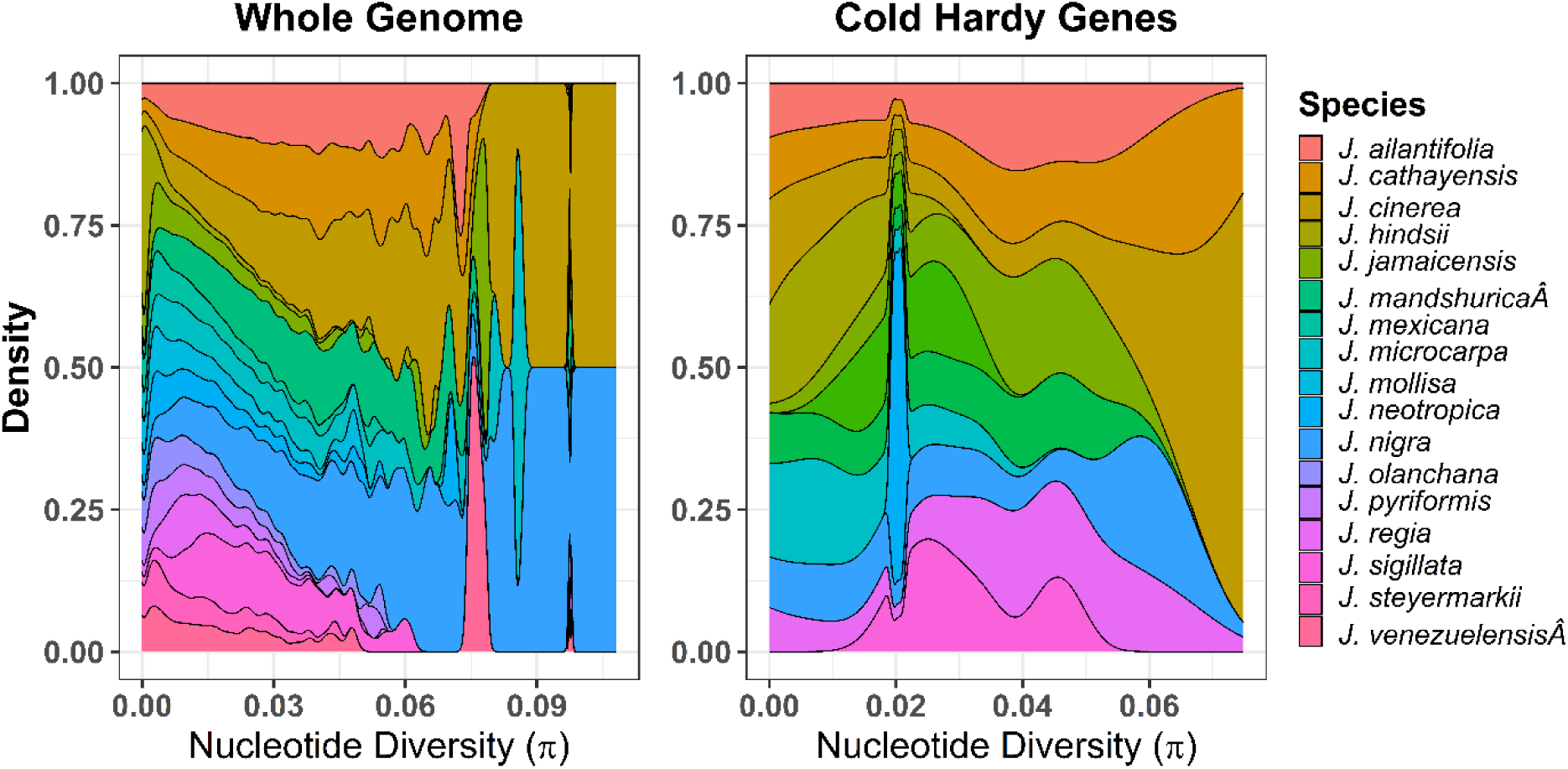
Nucleotide diversity of the whole genome (left) and cold-hardy genes (right) in the *Juglans* species.

